# Recognizability affects the processing of facial sex information

**DOI:** 10.1101/2025.06.26.661449

**Authors:** Szabolcs Sáringer, András Benyhe, Péter Kaposvári

**Affiliations:** Department of Physiology, Albert Szent-Györgyi Medical School, University of Szeged Dóm tér 10. 6720 Szeged, Hungary

**Keywords:** EEG, MVPA, face processing, familiarity, facial sex

## Abstract

The different aspects of a face, like sex or identity, can be decoded from the cortical patterns related to its processing. Many studies have investigated this phenomenon with similar outcomes. These studies usually utilize a low number of facial identities and high repetition numbers, which affects the recognizability and familiarity of a face, thus altering the processing. We propose that this commonly employed paradigm influences cortical patterns associated with features seemingly unrelated to identity, such as the sex of a face.

In the first experiment, we recreated the findings of previous studies using a few identities and a high presentation number for decoding facial sex. In the second experiment, the identity-presentation ratio was switched. This change resulted in a narrower time window where facial sex related cortical patterns were detected. Decoding accuracy was also diminished, yielding lower values and suggesting a reduced signal-to-noise ratio in the cortex. After expanding the sample size with balanced gender representation, we identified shared cortical patterns related to face-sex processing both within the population and across gender-based subpopulations. These results provide further evidence that familiarity impacts face processing and suggest that previous findings on sex information decoding were likely influenced by the experimental paradigms employed.

## Introduction

Fast and accurate face recognition is a crucial part of social life. It has been established that this process is tied to the hierarchical system of the ventral visual pathway (Axelrod & Yovel, 2012; Duchaine & Yovel, 2015; Zhen et al., 2013). Both behavioral (Barragan-Jason et al., 2013) and neural evidence (Goffaux & Dakin, 2010) suggest that different aspects of facial information processing are distributed across the ventral stream both spatially and temporally.

One such aspect is the sex of the observed face. Participants can discriminate between face sexes with high accuracy (Wild et al., 2000). Furthermore, behavioral data suggest a female advantage in face processing (Cellerino et al., 2004; Østergaard Knudsen et al., 2021), a possible hint that the mechanism might differ across genders (Proverbio et al., 2006). Besides the behavioral evidence, it was observed that face-sex information activates the core and extended face network (Kaul et al., 2011; Podrebarac et al., 2013) and follows its hierarchical structure. For example, previous studies show that face-sex information is processed prior to identity (Besson et al., 2017; Dobs et al., 2019), although some suggest the processes might run parallelly (Rossion, 2002).

It has also been established that the processing of familiar and unfamiliar faces differs in their neural signatures (Ambrus et al., 2019; Dobs et al., 2019; Kovács, 2020). Particularly, the degree of familiarity, or the experience leading to familiarity, can determine the cortical representation of a given face. The level of familiarity is greatly affected by the experienced visual variance in the face and the semantic information associated with said face. Based on this, previous studies implemented the familiarization process on different levels: mere visual familiarization provides small variance and no semantic information. In contrast, personal familiarization can result in greater effects (Ambrus et al., 2021). Recent papers also suggest that faces activate the same networks to different extents depending on the level of familiarity (Kovács, 2020).

Many studies have observed the underlying structure of the face-processing network extending from the primary visual cortex to the fusiform face area. A growing number of studies investigate the temporal dynamics of the process using M/EEG-based multivariate pattern analysis (MVPA) or representational similarity analysis (RSA) techniques (Ambrus et al., 2019, 2021; Dobs et al., 2019; Nemrodov et al., 2018).

MVPA and RSA analyses allow us to observe the temporal dynamics of several features simultaneously. Furthermore, these techniques also allow identifying shared cortical patterns across participants or experiments (Ambrus, 2024; Kaplan et al., 2015).

The results among the articles are consistent regarding early decoding differences, suggesting a short latency in the process. The different facial information (e.g., sex, identity, age) is coded in the cortical EEG pattern and can be distinguished as early as 50 ms, but generally speaking, most studies find a latency no longer than 100 ms. This is also consistent with behavioral results (short reaction times) and other imaging techniques (ECoG) where face-related neural signals are detectable; for example, the activation of face-selective areas in the inferotemporal cortex can be detected as early as 130 ms (Schrouff et al., 2020). A great disparity between the studies is the time window related to specific facial features and for how long the aspect-specific cortical patterns are maintained. Several of the above-mentioned studies found 500-800 ms long windows. A common feature among these studies is that they used a low number of facial identities with high repetition numbers and usually investigated several facial features simultaneously, like gender, identity, and familiarity. Although there is a great difference in the degree of familiarity, we can argue that mere exposure to unfamiliar faces will result in a familiarity effect (Ambrus et al., 2021). This experimental setup can familiarize the seemingly ‘unfamiliar’ facial stimuli during the experiment. Although the difference in the level of familiarity is undeniable in these studies, unfamiliar faces become recognizable using high repetition numbers.

A study, similar to the ones already mentioned, investigated the emergence of identity-specific cortical patterns in time. In contrast with many other publications, their goal only involved the observation of one facial feature (identity), and their methods included a high number of identities (91) with a relatively low number of repetitions (4) of one identity (Vida et al., 2017). Their findings were limited to a short time window (100 - 400 ms) despite the strong similarities with other RSA/MVPA studies. This raises the question whether subjects are familiarized with faces during the stimulus exposure and how this low level of familiarity affects the temporal dynamics of face processing.

Here, we argue that the previously unfamiliar faces gain familiarity and become recognizable through visual stimulation, even in experiments, and this familiarity affects facial features seemingly unrelated to familiarity, such as sex. To investigate the effect of visual exposure, we conducted two experiments. In Experiment 1, we set our goal to recreate the findings of previous MVPA/RSA studies with a high number of repeated displays (100×) of six identities and decode face-sex information from the related EEG activity. In Experiment 2, we switched the parameters by using 100 identities and a low number of repetitions (6×) for individual faces. The results of Experiment 2 indicate that reducing the recognizability of faces in a paradigm changes the temporal aspects of the cortical signal of face-sex processing.

Since the literature lacks similar paradigms, Experiment 2 was extended to further elaborate on face-sex processing. The balanced number of participants in the female and male gender subpopulations allows us to find similar or shared cortical patterns within the whole population, within a subpopulation, and between male and female subpopulations.

## Materials and Methods

### Experiment 1

#### Participants

In Experiment 1, 25 healthy volunteers (17 females, 23.85 (±1.26) years mean (±SD) age) with normal or corrected-to-normal vision participated. The sample size was determined based on previous publications employing similar methodologies. All participants gave written informed consent before registration, and the study protocol was approved by the Hungarian National Center for Public Health and Pharmacy (NNGYK/49248-3/2024).

#### Stimuli

One hundred stimuli were selected from the Chicago Face Database (Ma et al., 2015), balanced between sexes (50 males, 50 females, all identifying as white), with neutral facial expression. The images were centered, cropped into an oval shape, and presented in greyscale on a grey background. In Experiment 1, 6 faces (3 female and 3 male) were selected randomly for each participant.

The variance between the stimuli was evaluated using the Structural Similarity Index Measure (Wang et al., 2004). SSIM ranged between 0.89 and 0.94 between images, 0.91 on average (Fig. 1).

**Figure 1.**
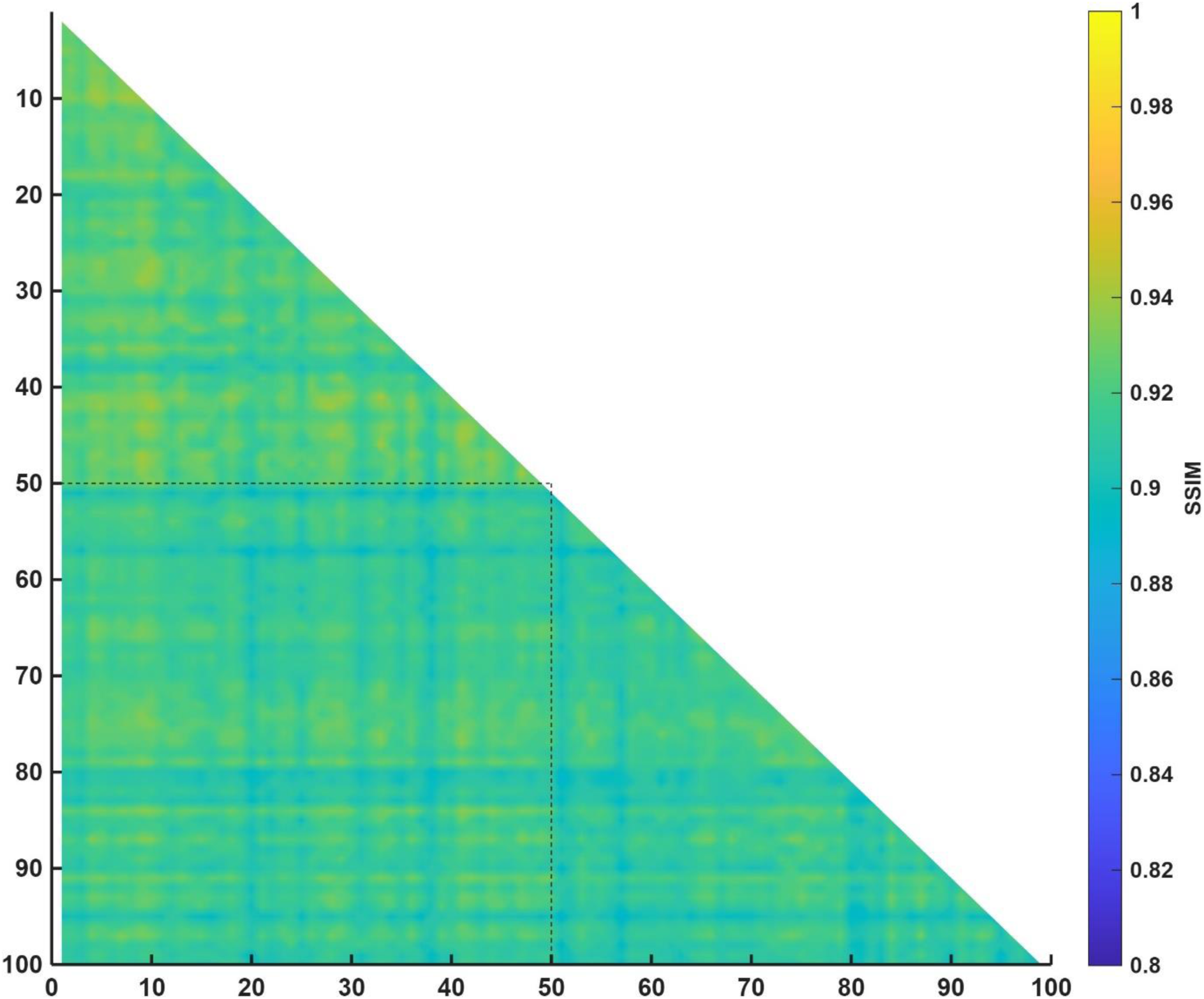
Structural Similarity Index Measure between stimuli. The first 50 images were of female faces, while the last 50 were of male faces. The average SSIM between pictures was above 0.9.

#### Procedure

Participants were seated approximately 50 cm from the screen and instructed that they were going to see a sequence of faces. The stimuli were presented using Psychtoolbox, MATLAB 2023 (Kleiner et al., 2007). They were also tasked with indicating the appearance of a target stimulus in the sequence with a button press. In target trials, the face stimulus was presented in a tilted manner (15° anticlockwise rotation). The purpose of the target stimuli was to maintain the participants’ attention throughout the image sequence. The EEG data of the target stimuli were excluded and not analyzed later.

All participants were presented with 600 stimuli (6 faces, 100 repetitions), of which 10% were target stimuli, thus resulting in 540 non-target stimuli for the subsequent analysis. A fixation cross was displayed at the center of the screen, which remained visible throughout the whole stimulus presentation. Stimuli (non-target and target) were presented for 500 ms, followed by a jittered intertrial interval of 800-1200 ms (Fig. 2). The whole sequence lasted approximately 15 minutes.

**Figure 2.**
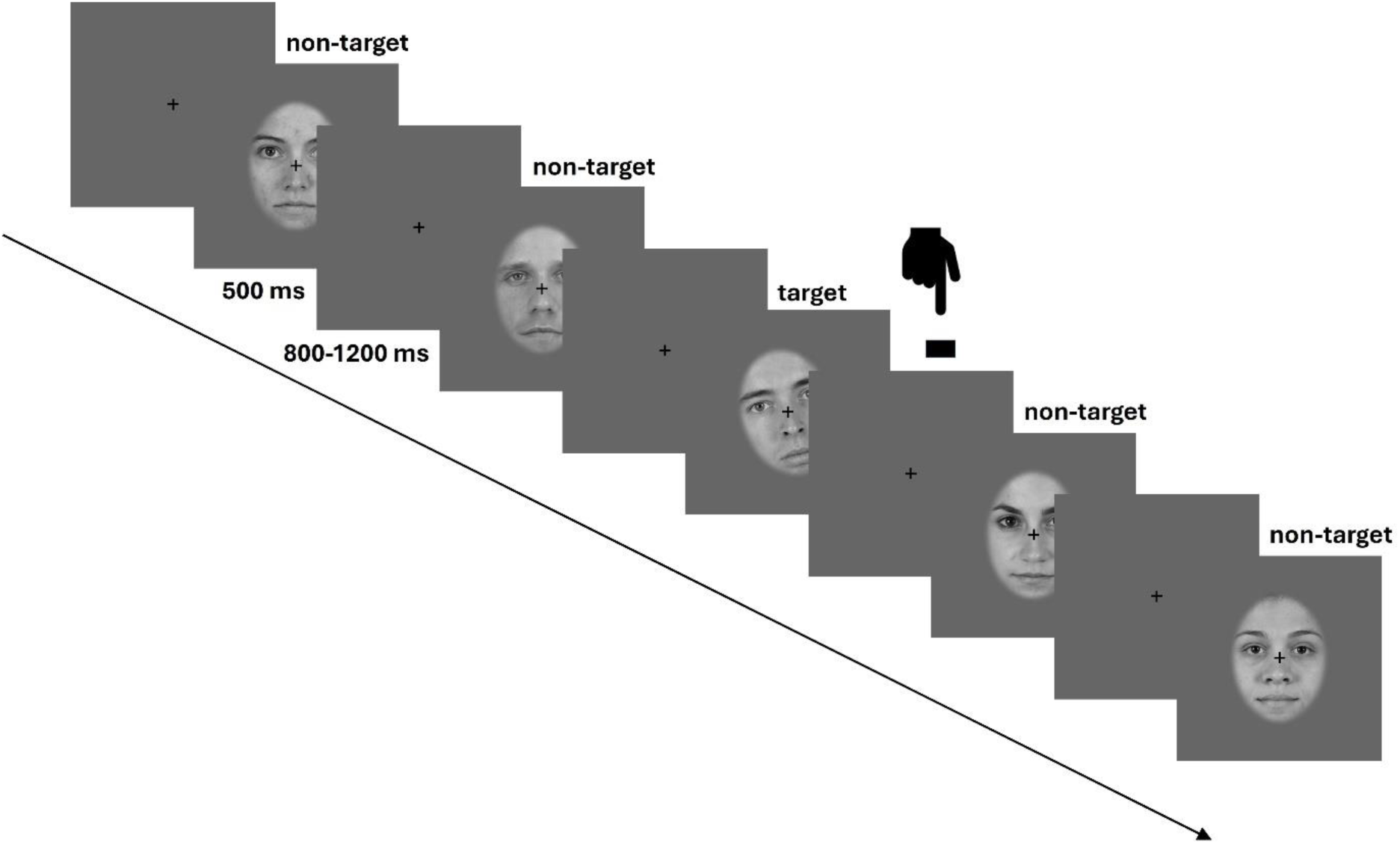
Stimulus presentation and structure

#### EEG acquisition and preprocessing

EEG data were recorded using a 64-channel Biosemi Active II System with a sampling rate of 512 Hz. Alongside the EEG, a 4-channel EOG was recorded from the outer canthi of the eyes and above and below from the left eye.

EEG preprocessing was implemented in MATLAB (2024a) using custom-written scripts and the FieldTrip toolbox (Oostenveld et al., 2010). First, a bandpass filter was applied to the raw data between 2 and 40 Hz. Then, eye-movement and blink artifacts were removed using a denoising source separation algorithm (Särelä & Valpola, 2005). We trained the algorithm on blink events using pseudotrials centered on EOG peaks. These components were then removed from the EEG data. Epochs were defined in the continuous data from 0.5 s before stimulus presentation to 1.2 s after. The data were resampled to 100 Hz and baselined to the prestimulus window of-0.3 to-0.1 s. Trials with unusually high variance were removed from the data. On average, 2.4 trials were removed per subject.

#### Data analysis

The behavioral results from the target detection task were used to determine subject exclusion. Subjects were to be excluded with a detection accuracy under 0.8, but all participants performed above this limit. To evaluate participants’ engagement in the task and experiment, mean accuracies and reaction times are reported.

To investigate the course of face-sex-specific cortical activity, we used Multivariate Pattern analysis (MVPA). Decoding accuracy was determined using the MVPA-Light and Fieldtrip toolboxes (Treder, 2020). A Linear Discriminant Analysis (LDA) algorithm was trained and tested on the face-sex of a given natural data at all time points of the first 1000 ms of the stimulus presentation. Cross-validation was performed with Leave-One-Out Cross-validation (LOOCV) method, where one trial was used as test data in one iteration. LOOCV was chosen due to the different identities and number of presentations between the experiments, and to keep the two results comparable. Similarly to previous related publications, a 60-ms-long moving mean window was applied to the decoding accuracy data (Vida et al., 2017).

Regional decoding was performed to acquire a limited spatial distribution of the activity. EEG electrodes were divided into six regions: left and right frontal, central, and occipital regions (following the methodology of Ambrus et al., 2019, 2021).

#### Statistics

The decoding accuracy of the algorithm was tested against chance (0.5) at all time points in the first second of the stimulus presentation using one-sample t-tests. Significant time windows were determined using Threshold-free Cluster Enhancement (TFCE) implemented in the Fieldtrip Toolbox. Correction due to multiple comparisons was performed against a null distribution created by permutations of 10000 iterations.

### Experiment 2

#### Participants

Experiment 2 included 25 participants (15 females, 25.70 (±4.44) years mean (±SD) age). All participants were healthy, with correct or corrected-to-normal vision, and agreed to participate in the experiment voluntarily. The experiment was accepted by the Hungarian National Center for Public Health and Pharmacy (NNGYK/49248-3/2024).

#### Procedure and data acquisition

The same stimulus bank, procedure, and data acquisition methods were used here as in Experiment 1. The difference in Experiment 2 is that we used all 100 stimuli and six repetitions, which reduced the familiarization process on a single face due to the low repetition number.

In Experiment 2, we used the same MVPA technique, with an LDA algorithm, and LOOCV. Decoding accuracies were evaluated, and significant time windows were determined in the same manner as in Experiment 1. Decoding accuracy across time was compared between Experiment 1 and Experiment 2 using independent samples t-tests with TFCE correction.

Additionally, across-experiment cross-validation was performed, where the training data came from the data of all subjects in one experiment, and the test set was set to be the data of one subject from the other experiment in one iteration. Just like previously, statistical significance was determined using independent t-tests and TFCE correction.

### Extension of Experiment 2

To keep the experiment balanced between genders and to raise statistical power, we expanded the number of subjects and recorded an additional 15 participants. This way, 40 volunteers participated in the extended experiment (20 females, 24.92 (±4.26) years mean (±SD) age). Different k-fold cross-validations were subsequently applied to the data to elaborate on the dynamics of face-sex decoding using a high number of identities and a high number of participants. The following cross-validations were applied in all possible permutations:

● *Within-subject CV:* Within-subject analysis included 99 identities in the training set, while the sex of 1 identity was tested using all repetitions of the face.
● *Between-subject CV:* To understand whether these cortical patterns are consistent between participants, we trained the LDA algorithm on the data of all participants except one, and the algorithm was tested on the excluded subject’s data.
● *Within-gender CV:* To investigate an assumed gender difference in face-processing based on behavioral data, our next analysis divided the population based on the gender of the participants. During within-gender analysis, the training set included the data of all participants of the subpopulation except one, which was used to test the algorithm’s accuracy.
● *Between-gender CV:* Analogous to the between-subject analysis, between-gender cross-validation was intended to test the general nature of the cortical

patterns between genders. The training set included the data of a subpopulation, while the algorithm was tested on the data of one participant in the subpopulation of the opposite gender.

### Results Experiment 1 Behavioral results

Accuracy in the detection task of the experiment was high, with a mean (±SD) of 0.97 (±0.03). The range was between 0.9 and 1. The average reaction time was 0.50 (±0.06) s.

#### LOOCV results

As expected, the decoding of face-sex information was successful in Experiment 1. An early time window can be observed between 85 ms and 475 ms, which is significantly above 0.5 (Fig. 3).

**Figure 3.**
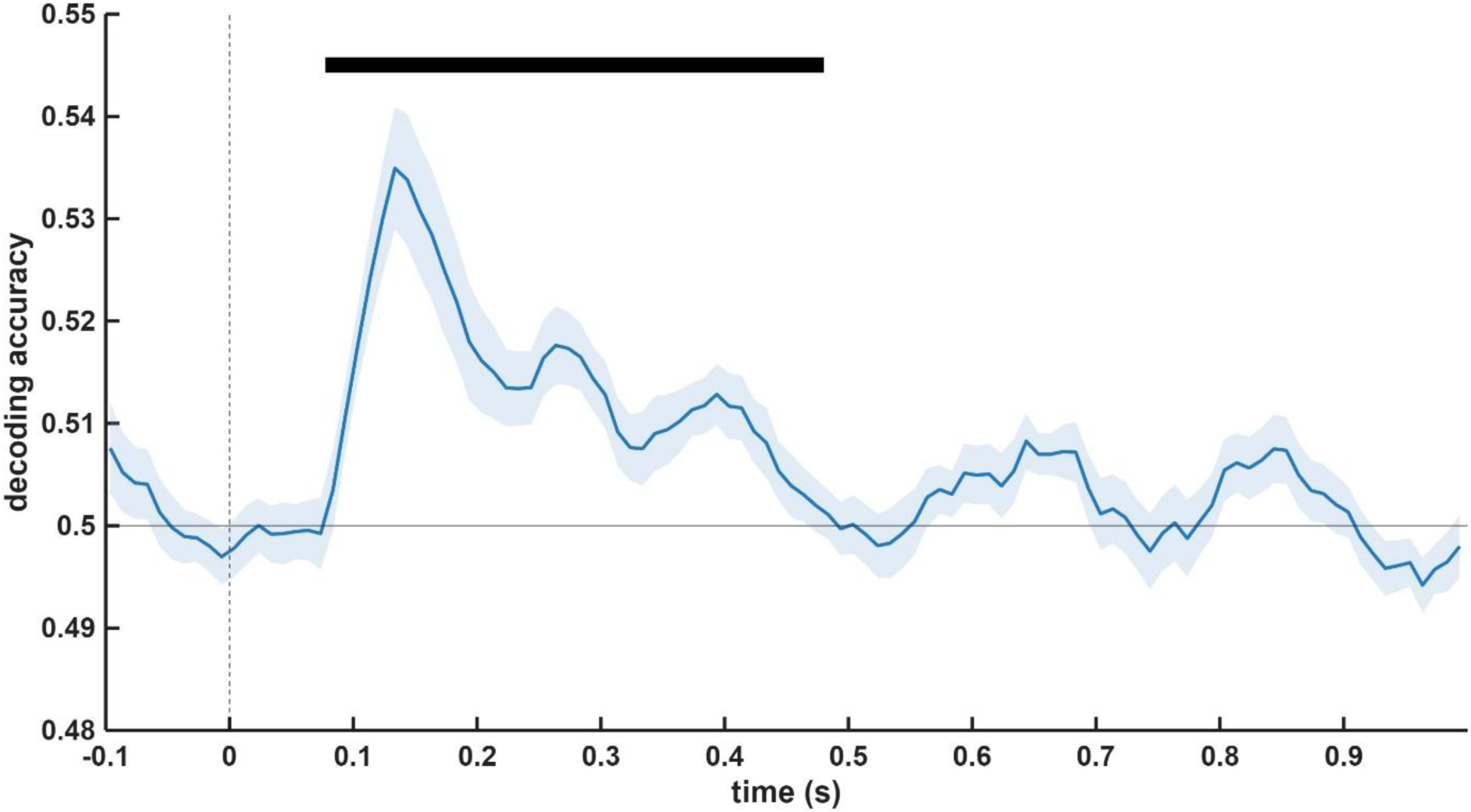
Decoding accuracy across time in Experiment 1. The shaded area marks SEM. Significant periods are indicated by a black bar.

Looking at the decoding accuracy in the previously defined electrode clusters revealed significantly above-chance time windows predominantly in the right-side regions. The greatest cluster can be observed in the occipital regions (left: 85-435 ms; right: 85-445 ms). Central regions also showed face-sex-related patterns, mainly on the right side (95-425 ms). Finally, shorter significant time windows also emerged in the right frontal regions between 195 ms and 885 ms (Fig. 4).

**Figure 4.**
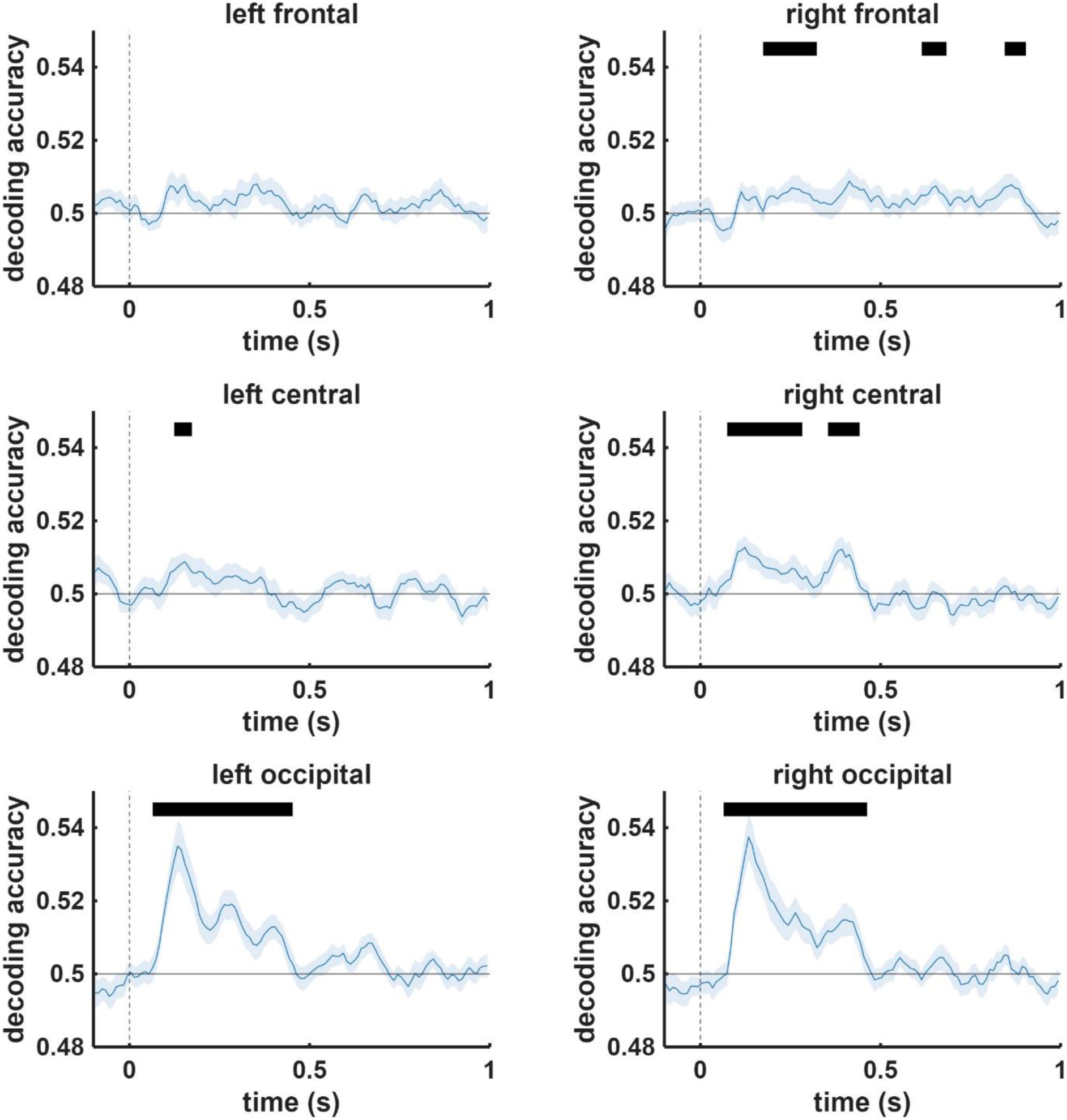
Decoding accuracy across time in Experiment 1 in the six regions described above. The shaded area marks SEM. Significant periods are indicated by a black bar.

### Experiment 2

#### Behavioral results

Similarly to Experiment 1, target detection accuracy was relatively high in Experiment 2 as well (mean (±SD) = 0.96 (±0.03), range: 0.88 - 1), and reaction time also showed comparable values (mean(±SD) = 0.48 (±0.05)).

#### LOOCV Results

It was also possible to decode the face-sex from the cortical patterns in Experiment 2, but the significant time window was shorter (95 to 285 ms) compared to the time window found in Experiment 1 (Fig. 5).

**Figure 5.**
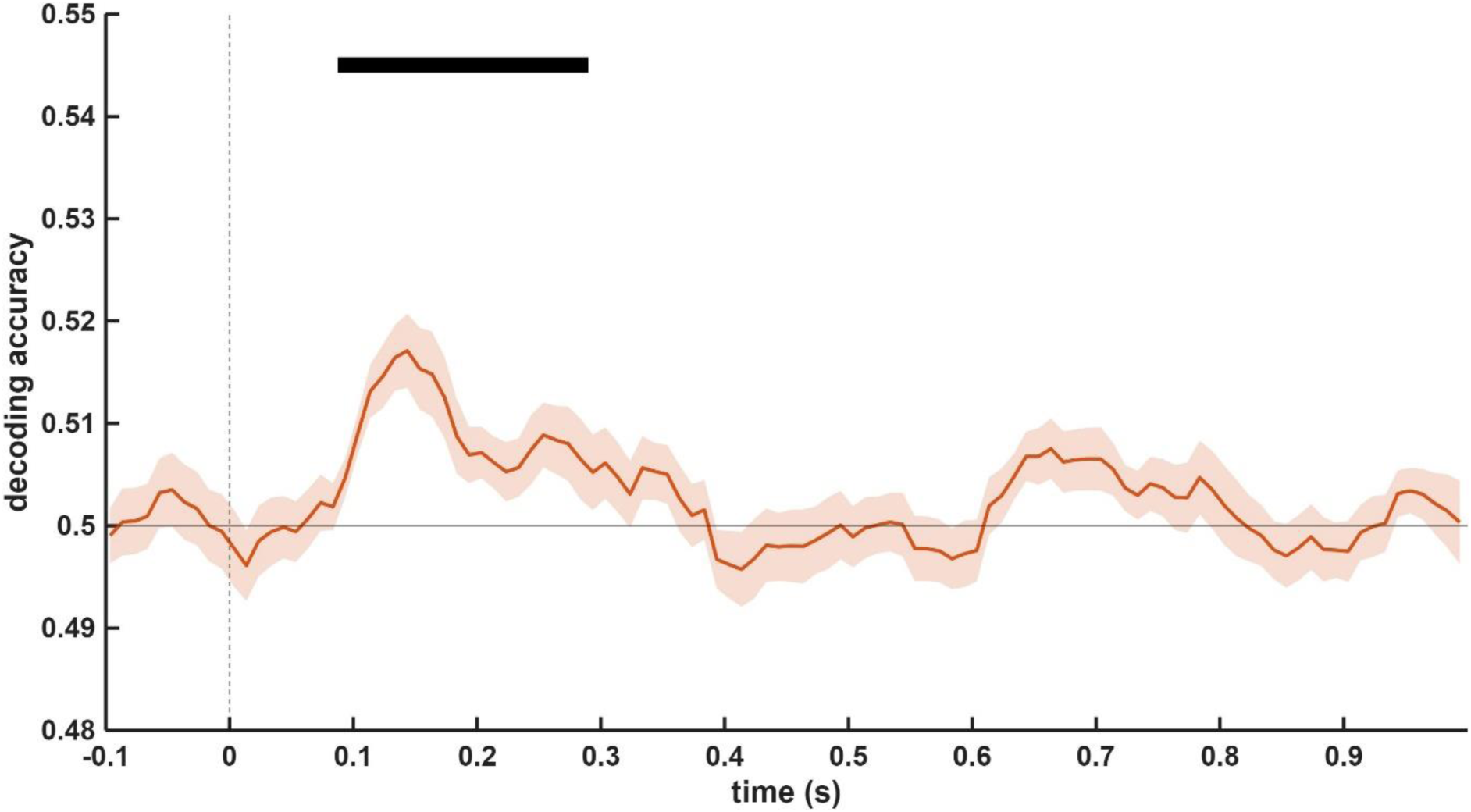
Decoding accuracy across time in Experiment 2. The shaded area marks SEM. Significant periods are indicated by a black bar.

Decoding in the different electrode clusters was also more restricted in Experiment 2. Significant clusters were only observed in the occipital regions with a similar shorter temporal distribution (left: 35-375 ms; right: 85-375 ms; Fig. 6).

**Figure 6.**
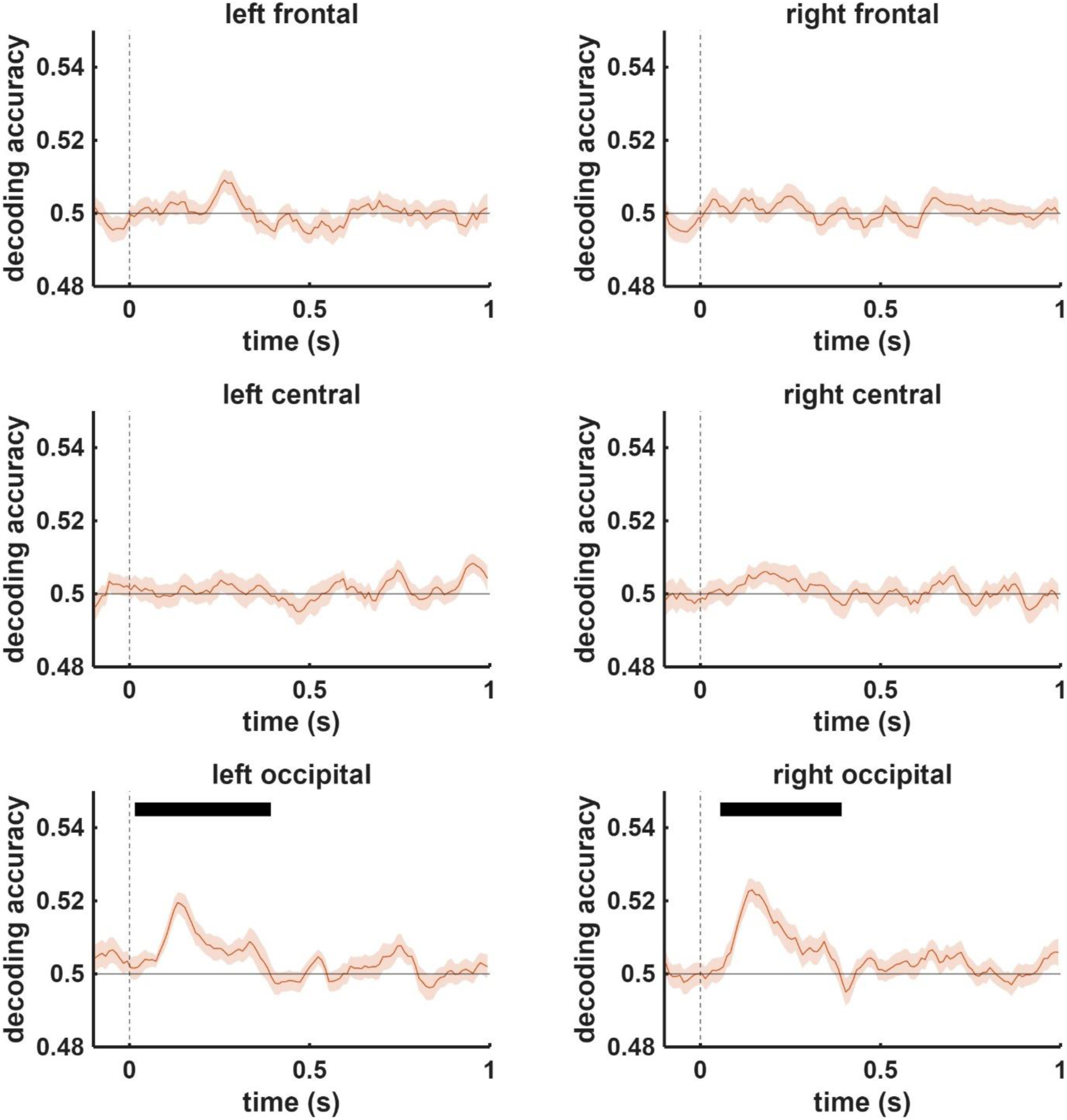
Decoding accuracy across time in Experiment 2 in the six regions described above. The shaded area marks SEM. Significant periods are indicated by a black bar.

### Comparison of Experiment 1 and Experiment 2

In the next step of the analysis, LDA accuracy was compared between the two experiments across time. This analysis revealed differences between 0.95 and 305 ms and 395-405 ms, where the algorithm performed significantly better in Experiment 1 (Fig. 7).

**Figure 7.**
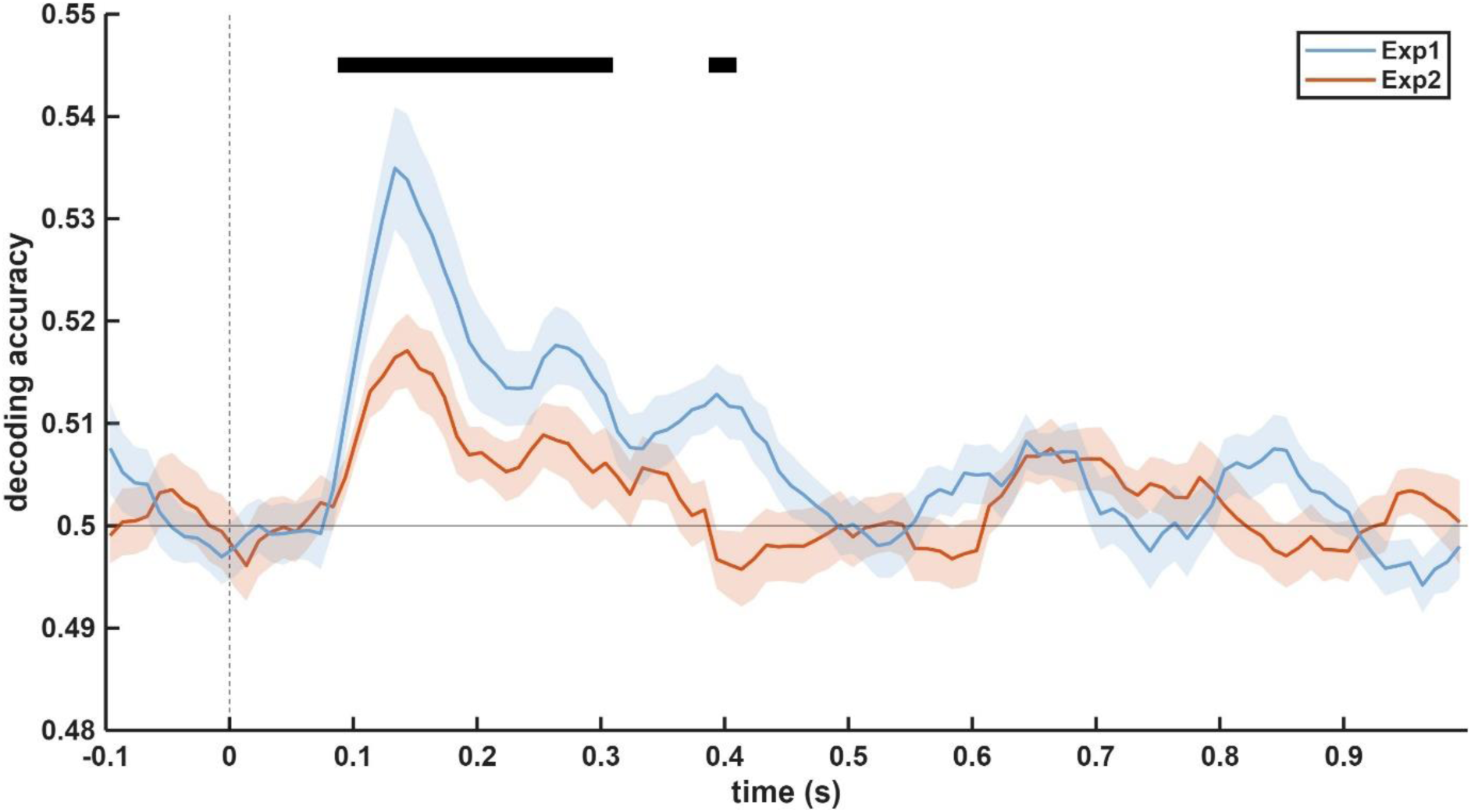
Comparison of the LDA performance in the two experiments across time. The blue line represents Experiment 1, while the orange line represents Experiment 2. The shaded area marks SEM, and the black bar signifies the significant time window.

Since the LOOCV results in an unbalanced representation of one identity (1 picture) between the two experiments, the data were reanalyzed with k-fold CV. In this case, in Experiment 1, one female and one male identity were left out of the training set and used as a testing set. Since only 6 identities were used in Experiment 1, one-third of the data was used as a testing set. To use the same training-testing ratio in Experiment 2, one-third of the trials were used as a testing set as well. This meant that through 3 iterations, 32-34-34 identities were left out (50% female, 50% male).

The reduction of the training set reduced the LDA performance in both experiments. In Experiment 1, a small, significant window appeared between 375 and 425 ms, while no window emerged in Experiment 2. The comparison of the two experiments resulted in no significant time window, either (Fig. 8).

**Figure 8.**
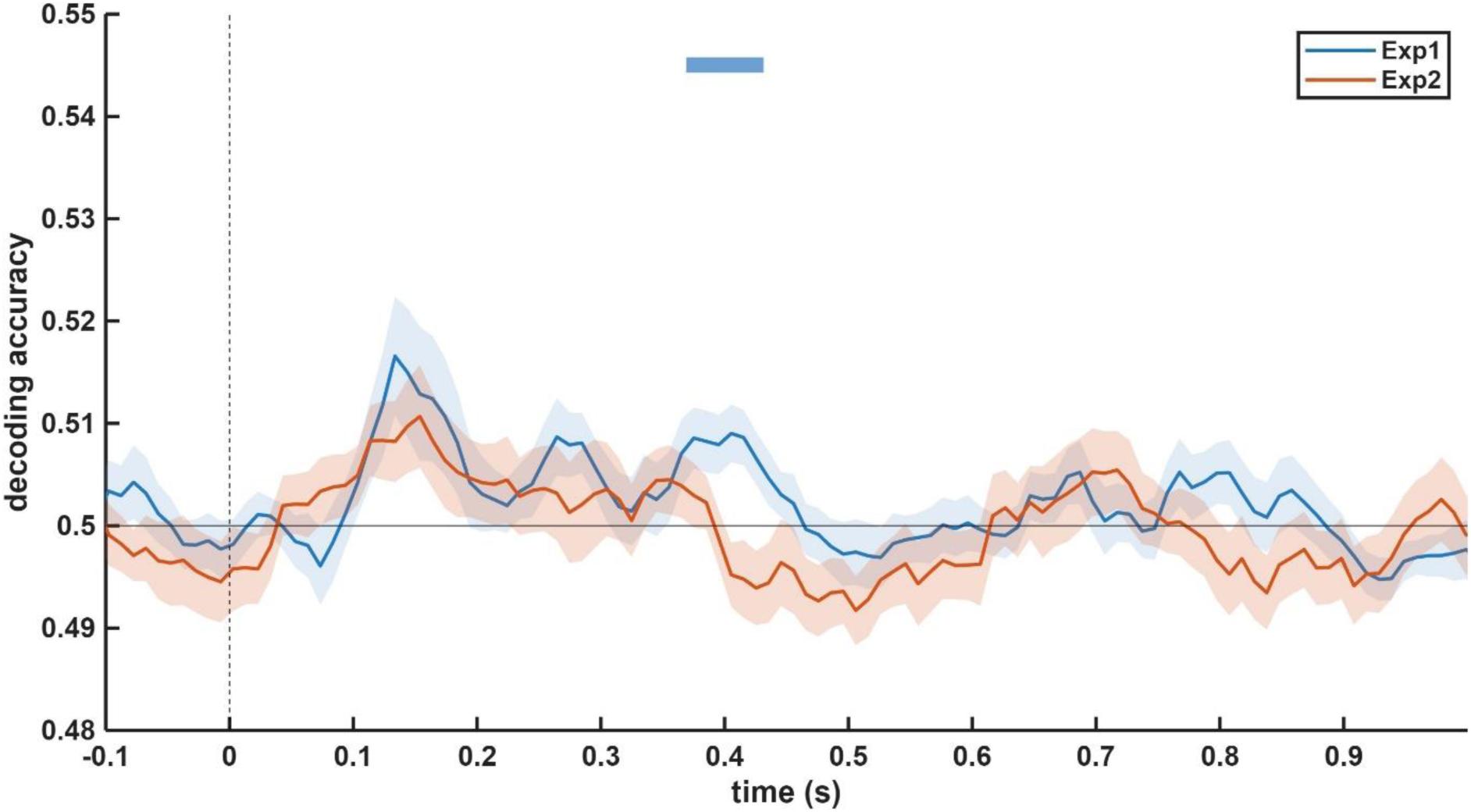
Comparison of the LDA performance in the two experiments across time using k-fold cross-validation. The blue line represents Experiment 1, while the orange line represents Experiment 2. The shaded area marks SEM, and the bar with the corresponding color represents the significant time window.

To further investigate the common neural patterns in face-sex decoding, we performed an across-experiment cross-validation. Training on Experiment 2 and testing on Experiment 1 revealed a significant window between 105 and 265 ms. Similarly, training on Experiment 1 and testing Experiment 2 resulted in an above-chance period between 105 and 255 ms (Fig. 9).

**Figure 9.**
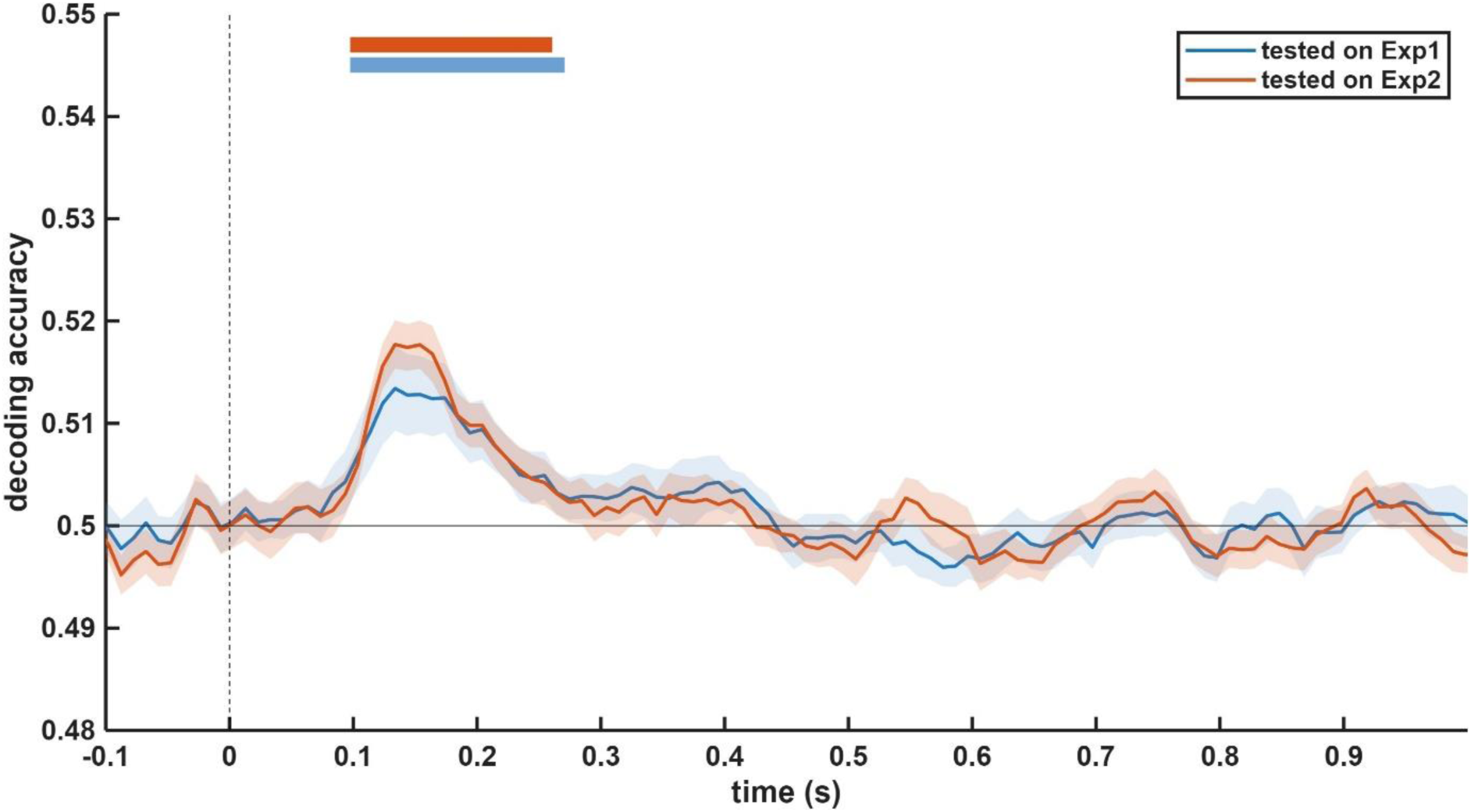
Decoding accuracy across time using across-experiment cross-validation. Significant periods are marked by the bars with corresponding colors. The data from Experiment 2 was used as a training set, while testing was performed on the data of one subject from Experiment 1 (blue line). The Training set was the data of Experiment 1, while the testing set was Experiment 2 (orange line).

### Extension of Experiment 2

Behavioral results remained high with the extension. Mean accuracy was above 0.8 for all subjects (mean (±SD) = 0.97 (±0.03), range: 0.88 - 1). Reaction time was fairly similar to the above-reported results (mean (±SD) = 0.49 (±0.05) s).

A within-subject CV, where one out of 100 identities was used for the k-fold cross-validation, revealed an early time window. Above-chance decoding was detectable between 75-355 ms post stimulus, unsurprisingly, almost identical to LOOCV analysis. Between-subject CV revealed a more restricted significant time window from 85 ms to 315 ms, characterized by an identical onset but a shorter duration compared to within-subject CV, along with an additional brief (20 ms) significant period around 740 ms (Fig. 10).

**Figure 10.**
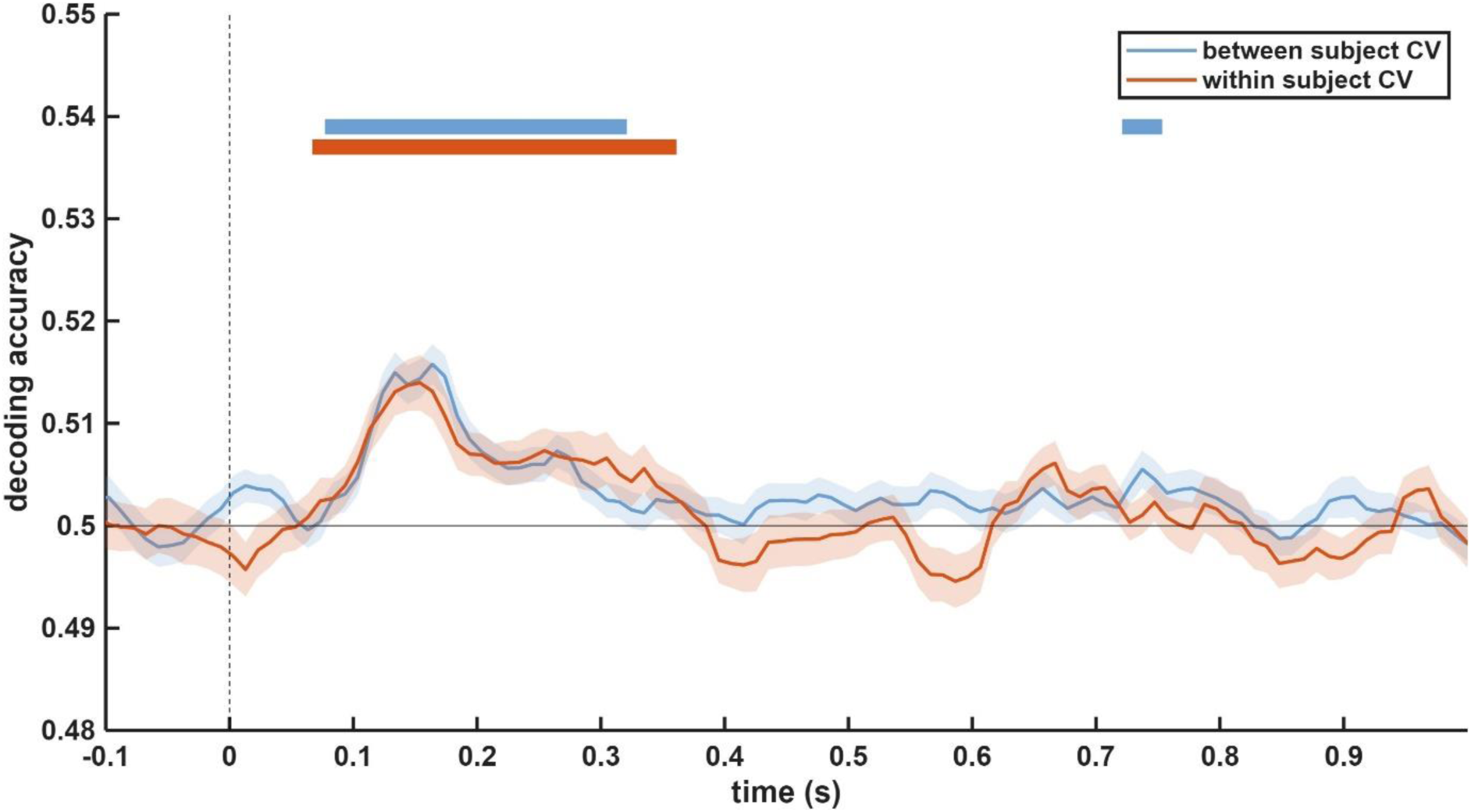
Decoding accuracy across time with within-subject CV and between-subject CV. The training set included all identities except one presented to one subject, which was subsequently used as a test set during within-subject CV (orange line). For the between-subject CV, the training set was determined as data from all subjects except one, which was used as a test set (blue line). Significant time windows are indicated by the corresponding bars.

Using within-gender CV revealed a temporal distribution for the above-chance period between 95-295 ms in the case of the male subpopulation and 115-275 ms in the case of the female subpopulation. This distribution is very similar to the between-subject CV, which can be assigned to the similarities between two cross-validations, as within-gender CV is fundamentally a between-subject CV within a subpopulation. Finally, the shortest significant time window was observed with between-gender CV, ranging from 115 ms to 205 ms when the model was trained on female data and tested on male data. When male data was used for training and tested on female data, no significant clusters emerged (Fig. 11).

**Figure 11.**
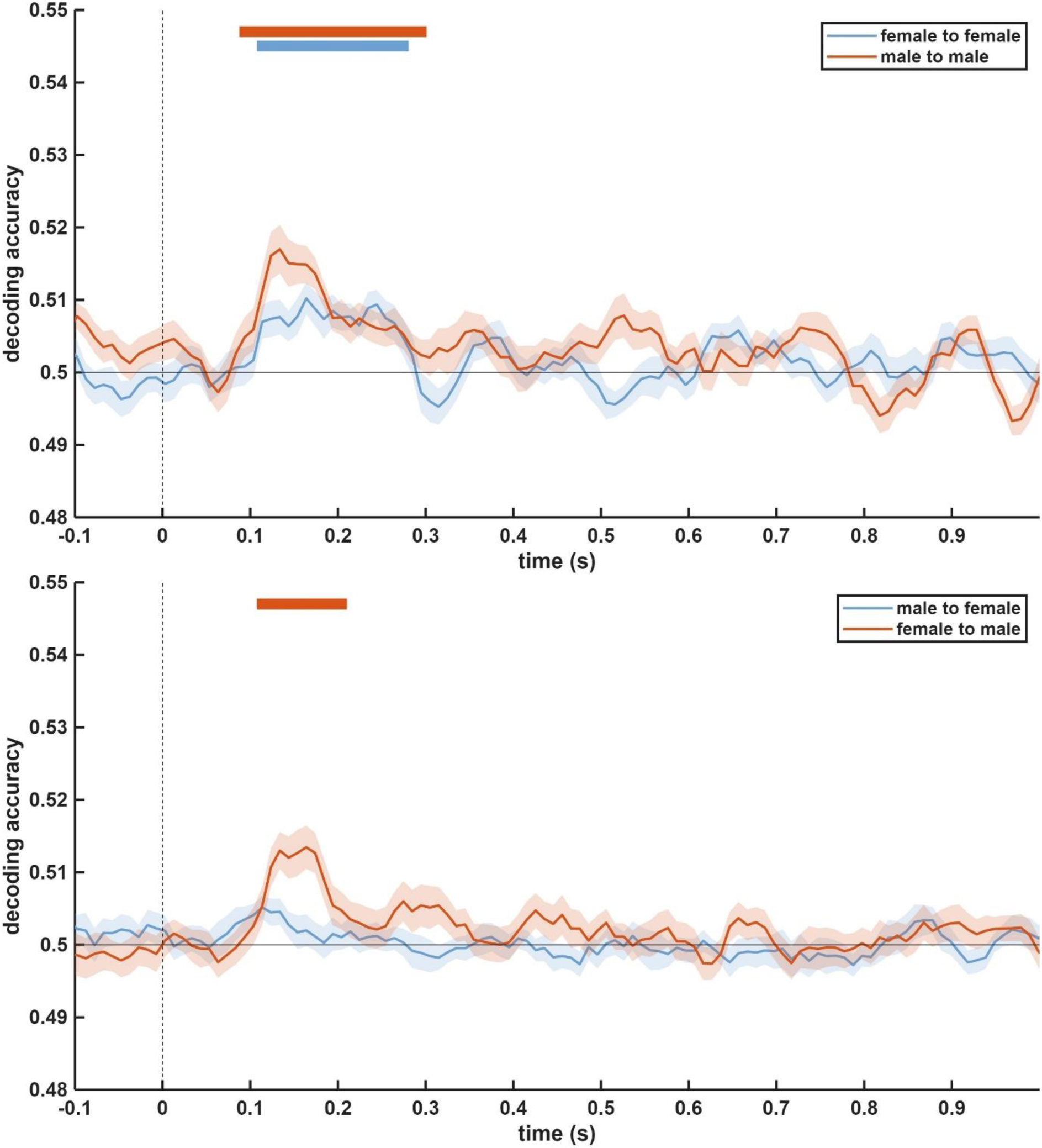
Top: Decoding accuracy across time using within-gender CV. The cross-validation used either the male or female subpopulation, where the data of all subjects except one was used as a training set and tested on one subject. The blue line represents the analysis where only the female subpopulation was used, while the orange line represents the male subpopulation. Bottom: Decoding accuracy with between-gender CV. The training set was determined as one subpopulation, either male or female. The test set was the data of one subject from the other subpopulation. The blue line represents the analysis where male subjects were used as a training set, and the model was tested on female participants. The orange line represents the opposite; the model was trained on the data of female participants and was tested on the data of male participants.

## Discussion

In this study, we investigated the effect of face recognizability and face familiarity on cortical decoding using different experimental setups. We hypothesized that previously reported MVPA/RSA results on facial sex decoding are influenced by the utilized paradigm. To test this hypothesis, we carried out two experiments where the degree of familiarity was altered using the number of presented identities and the number of presentations per identity.

In Experiment 1, we employed a paradigm commonly favored in previous research, characterized by a small number of identities coupled with high repetition numbers. Using these methods, we observed a time window where facial sex information can be decoded from the cortical pattern. The temporal distribution of this window was similar to the literature, with a very short latency of 80 ms and the latest significant periods as late as 880 ms. The regional analysis of the different cluster electrodes also showed a spatial distribution similar to previous reports, where identity information of familiar faces emerged over the occipital, central, and frontal regions, predominantly on the right side (Ambrus et al., 2019).

During Experiment 2, the ratio of identities and presentation numbers was switched, reducing the level of familiarity of one identity; other aspects of the stimulus presentation remained unchanged. This modification resulted in a more confined window where the sex of a facial stimulus can be decoded above chance. The distribution of the above-chance decoding was restrained not only temporally but spatially as well, since above-chance decoding was only detectable above the occipital regions. The comparison of the two experiments revealed that the identity/repetition ratio affects the decoding accuracy and thus the cortical patterns not only temporally and spatially but most likely in the signal-to-noise ratio as well, since the LDA algorithm performed significantly better in the early time windows in Experiment 1 compared to Experiment 2. Furthermore, with across-experiment cross-validation, we found a time window between 100 and 260 ms that appears to contain shared neural activity patterns between the two experiments.

After establishing the effect of the different paradigms, we extended the number of subjects in Experiment 2. With this extension, we were able to investigate the neural pattern and gender decoding of non-recognizable faces. We took the opportunity to use the high number of participants to find shared cortical patterns across subjects and the genders of observers. We identified significant time windows using different k-fold cross-validations, which suggests the existence of general or abstract patterns in the activation of the cortex, which are detectable by MVPA techniques.

### The decoding of facial sex information

Numerous publications have investigated the temporal distribution of cortical patterns related to face-sex processing. Previous studies have identified the time window when the sex of a presented face stimulus can be determined accurately. Most studies concluded that decoding accuracy emerges around 100 ms after stimulus presentation, with the earliest evidence found around 50 ms (Nemrodov et al., 2018). It is also shared between the studies that these decoding windows usually last about 600-700 ms, ending around 700-800 ms after the presentation of a stimulus. The regional analyses also revealed the occipital regions and the greater engagement of right central and frontal clusters (Ambrus et al., 2019). Besides the similarities in the results, many aspects of the applied paradigms are shared as well. Many studies use a low number of identities coupled with a lot of repetitions, for example, 16 identities and 140 repetitions per identity (Dobs et al., 2019), 4 identities and 400 repetitions per identity (Ambrus et al., 2019), 4 identities and 1300 repetitions per identity (Nemrodov et al., 2016). The use of high repetition numbers was inevitable in these studies since most investigated several facial features, like sex, identity, age, familiarity, etc.

Additionally, one identity included several stimuli/pictures, which boosted the variance in these paradigms. We already have evidence that changes in the neural pattern and decoding can emerge due to the perceptual familiarization process (Ambrus et al., 2021). Although perceptual familiarization has involved the smallest changes compared to media and personal familiarization processes, this effect cannot be neglected. Familiarity can be decoded from cortical activity, and the effect of familiarity on the face-processing mechanism has been established (Kovács, 2020).

So far, to our knowledge, only one other study has investigated identity-related cortical patterns using a high number of identities with a low repetition number (Vida et al., 2017). They reported a narrower significant window (100-400 ms), although the smaller sample size in their study might have contributed to this restricted timeframe. Our results align well with the literature, where perceptually familiarized faces induced sex-related cortical patterns to remain detectable for longer, while in the case of completely unfamiliar faces, these patterns are short-lived. Although many aforementioned studies investigated the decoding of familiarity, our findings provide evidence that the paradigms employed in those studies likely resulted in all presented faces acquiring some degree of familiarity. These results are also evidence that familiarity not only affects the processing as a whole, but it also affects seemingly unrelated features like the sex of a face.

We already have evidence that expectation can improve stimulus representation (Kok et al., 2012) and that familiar items can elicit superior perceptual performance (Manahova et al., 2018). The comparison of the two experiments also indicates that faces with a greater degree of familiarity elicit greater decoding accuracies, suggesting a better signal-to-noise ratio in the face-processing network. However, this difference is only detectable using LOOCV, which could result in overrepresentation of one identity during the training in Experiment 1. To investigate this, we reanalyzed the data with k-fold CV, where one-third of the trials were used for testing. This method drastically reduced the accuracy of the decoding algorithms in both experiments and removed the significant difference between the two. Nevertheless, significant decoding above chance was still achieved in Experiment 1, even with this reduced training set, suggesting a robust neural representation under these conditions, whereas Experiment 2 did not show significant decoding accuracies.

### Shared neural patterns in face-sex processing

There is a growing number of studies investigating the neural patterns associated with different aspects of perception and cognitive functions using decoding techniques. These studies usually use train and test sets within a subject, which is highly specific to personal cortical activity but does not provide any information about common patterns in the population. Recent studies started to examine these shared activities by using train and test sets across subjects or even experiments (Kaplan et al., 2015), and several cognitive functions have been identified that possess common cortical patterns, such as memory and recall (Ambrus, 2024), the reward system (Wake & Izuma, 2017), or familiarity (Dalski et al., 2022; C. Li et al., 2022).

Shared neural patterns in face perception have been identified as well. It has been established that face familiarity elicits shared activity 200 ms after stimulus presentation (Ambrus, 2024; Dalski et al., 2022). The investigation of the processing of different facial expressions and emotions has also led to the observation of shared neural patterns.

A previous study has investigated the common patterns in cortical activity regarding facial sex information and its perception (Y. Li et al., 2022). Their results showed that shared patterns could be detected in the EEG data around 150-400 ms after stimulus presentation. This finding is comparable to our between-subject CV results, which resulted in a 200-ms time window starting 100 ms after stimulus presentation. The difference in the length of these shared patterns could be attributed to differences in the paradigm, as in their case, 8 presented identities were used compared to the 100 used in this current study.

Our results confirmed and also expanded on these findings by using distinctly different subpopulations for training and testing. We already have behavioral and imaging data (Ino et al., 2010) that face perception differs between male and female participants. The current study found cortical patterns shared between the two subpopulations; however, these cross-validations yielded the shortest significant time windows, lasting about 80 ms only. The discrepancy found in the length of the significant time window, comparing within-gender and between-gender CV, can be a sign of gender specific cortical patterns. This difference and the lack of a significant time window for between-gender CV using female participants as a training set are a possible route to further explore gender differences in face processing.

## Conclusion

The current study investigated recognizability and its effect on maintaining facial sex information longer in the cortex. Our findings confirmed previous suspicions that using face perception paradigms with a small number of identities can greatly affect the processing of other facial aspects, in our case, the processing of sex information. By employing a large number of identities, we effectively reduced the degree of visual familiarity, which correlated with shorter significant decoding time windows and lower decoding accuracies, suggesting a diminished signal-to-noise ratio in the underlying cortical activity.

Furthermore, we extended the participants in the second part of the study to investigate the shared neural patterns in the population and between subpopulations (men and women). Both analyses identified significant time windows, suggesting the existence of common cortical patterns regarding facial sex processing, even between distinct subpopulations.

## Acknowledgement

We would like to thank our undergraduate student, Eszter Domboróczki, for her help in data acquisition and participant recruitment.

## Data availability

The data and scripts used for analyses in the current study are available in the Open Science Framework repository (https://osf.io/p593q/).

## CRediT statement

SzS: data curation, formal analysis, investigation, methodology, software, visualization, writing - original draft

AB: data curation, investigation, software, validation, writing - review & editing

PK: conceptualization, funding acquisition, methodology, project administration, supervision, validation, writing - review & editing

## References

Ambrus, G. G. (2024). Shared neural codes of recognition memory. Scientific Reports, 14(1), 15846. 10.1038/s41598-024-66158-y

Ambrus, G. G., Eick, C. M., Kaiser, D., & Kovács, G. (2021). Getting to Know You: Emerging Neural Representations during Face Familiarization. Journal of Neuroscience, 41(26), 5687–5698. 10.1523/JNEUROSCI.2466-20.2021

Ambrus, G. G., Kaiser, D., Cichy, R. M., & Kovács, G. (2019). The Neural Dynamics of Familiar Face Recognition. Cerebral Cortex, 29(11), 4775–4784. 10.1093/cercor/bhz010

Axelrod, V., & Yovel, G. (2012). Hierarchical processing of face viewpoint in human visual cortex. The Journal of Neuroscience, 32(7), 2442–2452. 10.1523/JNEUROSCI.4770-11.2012

Barragan-Jason, G., Besson, G., Ceccaldi, M., & Barbeau, E. J. (2013). Fast and Famous: Looking for the Fastest Speed at Which a Face Can be Recognized. Frontiers in Psychology, 4. 10.3389/fpsyg.2013.00100

Besson, G., Barragan-Jason, G., Thorpe, S. J., Fabre-Thorpe, M., Puma, S., Ceccaldi, M., & Barbeau, E. J. (2017). From face processing to face recognition: Comparing three different processing levels. Cognition, 158, 33–43. 10.1016/j.cognition.2016.10.004

Cellerino, A., Borghetti, D., & Sartucci, F. (2004). Sex differences in face gender recognition in humans. Brain Research Bulletin, 63(6), 443–449. 10.1016/j.brainresbull.2004.03.010

Dalski, A., Kovács, G., & Ambrus, G. G. (2022). Evidence for a General Neural Signature of Face Familiarity. Cerebral Cortex, 32(12), 2590–2601. 10.1093/cercor/bhab366

Dobs, K., Isik, L., Pantazis, D., & Kanwisher, N. (2019). How face perception unfolds over time. Nature Communications, 10(1), 1258. 10.1038/s41467-019-09239-1

Duchaine, B., & Yovel, G. (2015). A Revised Neural Framework for Face Processing. Annual Review of Vision Science, 1(Volume 1, 2015), 393–416. 10.1146/annurev-vision-082114-035518

Goffaux, V., & Dakin, S. C. (2010). Horizontal information drives the behavioral signatures of face processing. Frontiers in Psychology, 1, 143. 10.3389/fpsyg.2010.00143

Ino, T., Nakai, R., Azuma, T., Kimura, T., & Fukuyama, H. (2010). Gender Differences in Brain Activation During Encoding and Recognition of Male and Female Faces. Brain Imaging and Behavior, 4(1), 55–67. 10.1007/s11682-009-9085-0

Kaplan, J. T., Man, K., & Greening, S. G. (2015). Multivariate cross-classification: Applying machine learning techniques to characterize abstraction in neural representations. Frontiers in Human Neuroscience, 9. 10.3389/fnhum.2015.00151

Kaul, C., Rees, G., & Ishai, A. (2011). The Gender of Face Stimuli is Represented in Multiple Regions in the Human Brain. Frontiers in Human Neuroscience, 4. 10.3389/fnhum.2010.00238

Kleiner, M., Brainard, D., Pelli, D., Ingling, A., Murray, R., & Broussard, C. (2007). What’s new in psychtoolbox-3. Perception, 36(14), 1–16.

Kok, P., Jehee, J. F. M., & de Lange, F. P. (2012). Less is more: Expectation sharpens representations in the primary visual cortex. Neuron, 75(2), 265–270. 10.1016/j.neuron.2012.04.034

Kovács, G. (2020). Getting to Know Someone: Familiarity, Person Recognition, and Identification in the Human Brain. Journal of Cognitive Neuroscience, 32(12), 2205–2225. Journal of Cognitive Neuroscience. 10.1162/jocn_a_01627

Li, C., Burton, A. M., Ambrus, G. G., & Kovács, G. (2022). A neural measure of the degree of face familiarity. Cortex, 155, 1–12. 10.1016/j.cortex.2022.06.012

Li, Y., Zhang, M., Liu, S., & Luo, W. (2022). EEG decoding of multidimensional information from emotional faces. NeuroImage, 258, 119374. 10.1016/j.neuroimage.2022.119374

Ma, D. S., Correll, J., & Wittenbrink, B. (2015). The Chicago face database: A free stimulus set of faces and norming data. Behavior Research Methods, 47(4), 1122–1135. 10.3758/s13428-014-0532-5

Manahova, M. E., Mostert, P., Kok, P., Schoffelen, J.-M., & de Lange, F. P. (2018). Stimulus Familiarity and Expectation Jointly Modulate Neural Activity in the Visual Ventral Stream. Journal of Cognitive Neuroscience, 30(9), 1366–1377. 10.1162/jocn_a_01281

Nemrodov, D., Niemeier, M., Mok, J. N. Y., & Nestor, A. (2016). The time course of individual face recognition: A pattern analysis of ERP signals. NeuroImage, 132, 469–476. 10.1016/j.neuroimage.2016.03.006

Nemrodov, D., Niemeier, M., Patel, A., & Nestor, A. (2018). The Neural Dynamics of Facial Identity Processing: Insights from EEG-Based Pattern Analysis and Image Reconstruction. eNeuro, 5(1). 10.1523/ENEURO.0358-17.2018

Oostenveld, R., Fries, P., Maris, E., & Schoffelen, J.-M. (2010). FieldTrip: Open Source Software for Advanced Analysis of MEG, EEG, and Invasive Electrophysiological Data. Computational Intelligence and Neuroscience, 2011, e156869. 10.1155/2011/156869

Østergaard Knudsen, C., Winther Rasmussen, Katrine, & and Gerlach, C. (2021). Gender differences in face recognition: The role of holistic processing. Visual Cognition, 29(6), 379–385. 10.1080/13506285.2021.1930312

Podrebarac, S. K., Goodale, M. A., van der Zwan, R., & Snow, J. C. (2013). Gender-selective neural populations: Evidence from event-related fMRI repetition suppression. Experimental Brain Research, 226(2), 241–252. 10.1007/s00221-013-3429-0

Proverbio, A. M., Brignone, V., Matarazzo, S., Del Zotto, M., & Zani, A. (2006). Gender differences in hemispheric asymmetry for face processing. BMC Neuroscience, 7(1), 44. 10.1186/1471-2202-7-44

Rossion, B. (2002). Is sex categorization from faces really parallel to face recognition? Visual Cognition, 9(8), 1003–1020. 10.1080/13506280143000485

Särelä, J., & Valpola, H. (2005). Denoising Source Separation. The Journal of Machine Learning Research, 6, 233–272.

Schrouff, J., Raccah, O., Baek, S., Rangarajan, V., Salehi, S., Mourão-Miranda, J., Helili, Z., Daitch, A. L., & Parvizi, J. (2020). Fast temporal dynamics and causal relevance of face processing in the human temporal cortex. Nature Communications, 11(1), 656. 10.1038/s41467-020-14432-8

Treder, M. S. (2020). MVPA-Light: A Classification and Regression Toolbox for Multi-Dimensional Data. Frontiers in Neuroscience, 14. 10.3389/fnins.2020.00289

Vida, M. D., Nestor, A., Plaut, D. C., & Behrmann, M. (2017). Spatiotemporal dynamics of similarity-based neural representations of facial identity. Proceedings of the National Academy of Sciences of the United States of America, 114(2), 388–393. 10.1073/pnas.1614763114

Wake, S. J., & Izuma, K. (2017). A common neural code for social and monetary rewards in the human striatum. Social Cognitive and Affective Neuroscience, 12(10), 1558–1564. 10.1093/scan/nsx092

Wang, Z., Bovik, A. C., Sheikh, H. R., & Simoncelli, E. P. (2004). Image quality assessment: From error visibility to structural similarity. IEEE Transactions on Image Processing, 13(4), 600–612. 10.1109/TIP.2003.819861

Wild, H. A., Barrett, S. E., Spence, M. J., O’Toole, A. J., Cheng, Y. D., & Brooke, J. (2000). Recognition and Sex Categorization of Adults’ and Children’s Faces: Examining Performance in the Absence of Sex-Stereotyped Cues. Journal of Experimental Child Psychology, 77(4), 269–291. 10.1006/jecp.1999.2554

Zhen, Z., Fang, H., & Liu, J. (2013). The Hierarchical Brain Network for Face Recognition. PLOS ONE, 8(3), e59886. 10.1371/journal.pone.0059886

